# FLYNC: A Machine Learning-Driven Framework for Discovering Long Non-Coding RNAs in *Drosophila melanogaster*

**DOI:** 10.1101/2024.02.27.582305

**Authors:** Ricardo F. dos Santos, Tiago Baptista, Graça S. Marques, Catarina C. F. Homem

## Abstract

Non-coding RNAs have increasingly recognized roles in critical molecular mechanisms of disease. However, the non-coding genome of *Drosophila melanogaster*, one of the most powerful disease model organisms, has been understudied. Here, we present FLYNC – FLY Non-Coding discovery and classification – a novel machine learning-based model that predicts the probability of a newly identified RNA transcript being a long non-coding RNA (lncRNA). Integrated into an end-to-end bioinformatics pipeline capable of processing single-cell or bulk RNA sequencing data, FLYNC outputs potential new non-coding RNA genes. FLYNC leverages large-scale genomic and transcriptomic datasets to identify patterns and features that distinguish non-coding genes from protein-coding genes, thereby facilitating lncRNA prediction. We demonstrate the application of FLYNC to publicly available *Drosophila* adult head bulk transcriptome and single-cell transcriptomic data from *Drosophila* neural stem cell lineages and identify several novel tissue- and cell-specific lncRNAs. We have further experimentally validated the existence of a set of FLYNC positive hits by qPCR. Overall, our findings demonstrate that FLYNC serves as a robust tool for identifying lncRNAs in *Drosophila melanogaster*, transcending current limitations in ncRNA identification and harnessing the potential of machine learning.

## INTRODUCTION

Traditionally, the study of stem cell function during animal development or tissue maintenance has usually emphasized the protein-coding genome. However, recent transcriptomic analyses have revealed that while coding genes constitute a small fraction of the genome, a significantly larger portion of the genome is transcribed(1). This extensive transcription of non-coding genome regions has been observed across various organisms, ranging from *Drosophila* to human(2, 3). Despite the wealth of knowledge derived from studying protein-coding genes (PCGs), research on non-coding genome has however lagged(4), likely due to the technical challenges associated with identifying and characterizing the function of non-conding RNAs (ncRNAs) *in vivo*. Non-coding regions encompass a diverse array of transcripts, with the most prevalent being long non-coding RNAs (lncRNAs). lncRNAs are defined as RNAs longer than 200 nucleotides that do not encode functional proteins(5). lncRNAs have been identified as regulators of various cellular processes, including chromatin dynamics, transcriptional regulation, translation control, pluripotency, and differentiation(6). Nevertheless, the exact *in vivo* functions of lncRNAs remain unclear, since its molecular functions have primarily been explored *in vitro*(7–13).

In fact, the study of individual lncRNAs poses significant challenges in vertebrates due to the intricate and time-consuming process of mutant generation. In contrast, *Drosophila melanogaster* emerges as an excellent model system for *in vivo* studies elucidating the functional roles of lncRNAs. Its genetic tractability, short generation time, well-characterized developmental processes, and conservation of many developmental pathways with mammals collectively render it an ideal platform for investigating complex biological phenomena. Also, brain development is a particularly intricate process involving the generation of thousands of different types of neurons and glial cells from a relatively small number of neural stem cells (NSCs). Pertinently, a large fraction of lncRNAs were found to be expressed specifically in the brain, in a cell-type and developmental stage-specific manner, correlating with the complex requirement for spatio-temporal gene regulation during brain development and neural differentiation(14). Once again, D. *melanogaster* NSCs and their lineages are extensively described, and the availability of powerful genetic tools enables precise manipulation of gene and lncRNA expression. However, despite these advantages, the non-coding genome of D. *melanogaster* remains poorly characterized, hindering the systematic utilization of this model for studying the *in vivo* function of lncRNAs. Therefore, there is an urgent need for a comprehensive characterization of *Drosophila*’s non-coding genome and the identification of lncRNAs. Such efforts will facilitate the systematic and integrated study of lncRNA function *in vivo* during animal development.

As lncRNAs exhibit weak conservation at the genomic sequence level, relying solely on genomic sequence for their identification poses significant challenges(15). Also, considering the limited understanding of the sequence-structure-function relationship of lncRNAs, their identification has primarily relied on transcriptome studies, namely via the advent of high-throughput RNA-sequencing technology(16). However, the usage of bulk transcriptome data limits the identification of cell-type-specific lncRNAs, despite their demonstrated regulatory roles and greater tissue specificity compared to mRNAs(17). Furthermore, the relatively low expression levels of most lncRNAs pose challenges in their detection, particularly in weakly sampled or broad transcriptomic datasets(18). Addressing the lack of cell specificity implies the analysis of single-cell transcriptomic datasets. With the emergence of platforms like 10x Genomics, numerous single-cell datasets are now available for analysis in *Drosophila*, spanning various tissues, organs, developmental stages, and mutant backgrounds(19, 20). These datasets offer unprecedented opportunities for precise identification of cell-specific non-coding regions in *Drosophila*, enabling comprehensive investigations into the regulatory roles and functional significance of lncRNAs throughout development and in diverse biological contexts.

However, accurate automated or manual annotation of lncRNA genes remains a challenging endeavor. Automated annotation relies on transcriptome assembly approaches, which are efficient and cost-effective but often yield incomplete and inaccurate annotations, leading to numerous false positives. On the other hand, manual annotation, exemplified by databases like RefSeq or GENCODE, involves rigorous experimental validation resulting in high-quality data, while being resource-intensive and time-consuming(16). Machine learning-based approaches have shown great promise in addressing various biological questions(21). These approaches can leverage large-scale genomic and transcriptomic data to identify patterns and features that distinguish ncRNAs from PCGs, thereby facilitating the prediction of functional ncRNAs in newly identified RNA transcripts(22). The integration of machine learning techniques with biological data has the potential to uncover novel ncRNA functions and contribute to a more comprehensive understanding of gene regulation.

In this paper, we introduce a novel machine learning-based model – FLYNC – designed to predict the likelihood of a newly identified RNA transcript being a ncRNA rather than a PCG in *Drosophila melanogaster*. Our model forms an integral part of an end-to-end bioinformatics pipeline that processes sequencing data and generates potential new ncRNA genes. By addressing existing limitations in bioinformatics tools for ncRNA research and leveraging the capabilities of machine learning, our model aims to enhance our understanding of the functional roles of ncRNAs in *Drosophila* development and broader biological contexts.

## MATERIALS AND METHODS

### Data sources

For the development of FLYNC, we integrated data from three reputable and widely recognized sources, each serving a distinct purpose and contributing unique datasets necessary for the comprehensive analysis of *Drosophila melanogaster* genomics data.

#### Sequence Read Archive (SRA)

The Sequence Read Archive (ncbi.nlm.nih.gov/sra) was selected as the primary source for sequencing reads due to its extensive repository of high-throughput sequencing data. The SRA is managed by the National Center for Biotechnology Information (NCBI) and is part of the International Nucleotide Sequence Database Collaboration (INSDC). It provides a platform for the storage and access of sequence data from a multitude of species and is instrumental in ensuring that the sequencing data used in our pipeline are both current and comprehensive. By leveraging the SRA’s remote access capabilities, our pipeline can directly download specific datasets on demand, ensuring efficient data management and up-to-date information for analysis.

#### Ensembl

Ensembl (ensembl.org) was chosen for sourcing reference genome annotations of *Drosophila melanogaster.* Ensembl is a project that provides a centralized resource for researchers seeking genome annotations. It is known for its high-quality, manually curated annotation data. The use of Ensembl annotations in our pipeline allows for the precise identification of genomic elements, such as genes, exons, and regulatory regions.

#### University of California, Santa Cruz (UCSC)

The University of California, Santa Cruz (UCSC) Genome Browser (genome.ucsc.edu) was utilized to extract genome features necessary for the construction of our machine learning models. The UCSC Genome Browser is a sophisticated and interactive tool that provides a user-friendly interface for visualizing genomic data and extracting features such as chromatin states, transcription factor binding sites, expression profiles, and comparative genomics information. The integration of UCSC data into our pipeline provides a rich feature set that enhances the FLYNC ML classifier’s ability to identify patterns and make predictions based on the genomic context. The comprehensive data available through UCSC are invaluable for the development of a robust and predictive bioinformatics pipeline.

### Computational resources

#### Specs of Workstation

For the development and testing of FLYNC we used Intel® Core™ i7-9700 computer with 64GB DRAM and NVIDIA Quadro P2000 GPU with 5GB GDDR5. This workstation has been used throughout the development of FLYNC and SUBCELL.

#### HPC information & Acknowledgements

The HPC service from the Lisbon node (cirrus.a.incd.pt) of Infrasturutra Nacional de Computação Distribuída (INCD) running CentOS 7 was mainly used for compute- and/or memory-intensive tasks, such as read alignment, transcriptome assembly, ML model training & evaluation.

### Training dataset and machine learning

To explore our evidence-based approach for features that would help distinguish lncRNAs from other genes in *D. melanogaster*, a pre-training sample dataset with a balanced representation NCG and PCG (10000 randomly selected genes) was prepared. This sample dataset was used to quickly assess suitable ML algorithms using ‘lazypredict’ Python library.

The selection of ‘scikit-learn’ as the machine learning framework for FLYNC was informed by several considerations. Python’s prominence in the data science community, coupled with its comprehensive ecosystem of libraries, makes it an ideal choice for the development of data-driven applications. The focus on supervised learning algorithms within ‘scikit-learn’ aligns perfectly with the classification nature of our task, which involves predicting lncRNA transcripts. Moreover, the framework’s emphasis on explainable models is particularly advantageous, as it allows researchers to interpret the biological significance of the features contributing to the predictions. Additionally, ‘scikit-learn’ offers numerous benefits, including a consistent API, comprehensive documentation, and a supportive community, making it an efficient tool for both prototyping and production.

The training dataset itself was crafted to incorporate features that are biologically relevant for the discrimination between non-coding and coding RNA transcripts. This carefully selected combination of features was chosen to reflect the functional importance and coding potential of RNA transcripts. The complete training dataset is provided available in FLYNC code repository, Detailed descriptions of these features and the rationale behind their selection are discussed in the “RESULTS & DISCUSSION” section.

### Evaluation of ML classifier

To objectively assess the performance of the Random Forest (RF) model used in FLYNC, we employed a suite of standard evaluation metrics. These metrics provide a comprehensive picture of the model’s predictive accuracy, its ability to distinguish between classes, and the balance between sensitivity and specificity. Below, we describe the evaluation metrics and the formulas used to calculate them.

#### Accuracy

Accuracy is the most intuitive performance measure, and it is simply a ratio of correctly predicted observation to the total observations. It is a useful metric only when the classes are balanced.

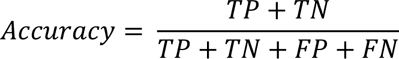

Where: TP = True Positives; TN = True Negatives; FP = False Positives; FN = False Negatives

#### Precision (Positive Predictive Value)

Precision is the ratio of correctly predicted positive observations to the total predicted positive observations. High precision relates to the low false positive rate.

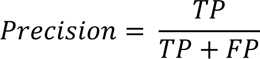

#### Recall (Sensitivity, True Positive Rate)

Recall is the ratio of correctly predicted positive observations to the all observations in the actual class. It is also known as Sensitivity or the True Positive Rate (TPR).

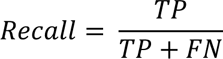

#### F1 Score

The F1 Score is the weighted average of Precision and Recall. Therefore, this score takes both false positives and false negatives into account.

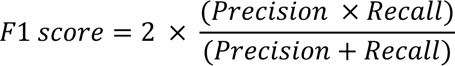

#### Confusion Matrix

A confusion matrix is a table that is often used to describe the performance of a classification model on a set of test data for which the true values are known. It allows the visualization of the performance of an algorithm.

#### Precision-Recall Curve

The precision-recall curve shows the trade-off between precision and recall for different thresholds. A high area under the curve represents both high recall and high precision.

#### Receiver Operating Characteristic (ROC) Curve

The ROC curve is a graphical plot that illustrates the diagnostic ability of a binary classifier system as its discrimination threshold is varied. It is created by plotting the True Positive Rate (TPR) against the False Positive Rate (FPR) at various threshold settings.

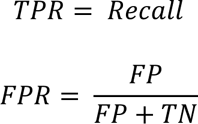

Where: TPR = True Positive Rate; FPR = False Positive Rate

#### Cross-Validation

To validate the performance of the RF model, we used 5-fold cross-validation for all metrics. This method involves randomly dividing the set of observations into five groups, or folds, of approximately equal size. The first fold is treated as a validation set, and the method is fit on the remaining four folds. This process is repeated five times, with each fold used exactly once as the validation data. This approach helps in mitigating overfitting and provides a robust estimate of the model’s performance on unseen data.

The performance metrics were computed for each of the five folds, and the results were averaged to provide a final performance estimate. The standard deviation of the performance across the folds was also calculated to assess the variability of the model’s performance.

### FLYNC CLI

The FLYNC (Fly Non-Coding RNA discovery and classification) Command Line Interface (CLI) is a powerful tool implemented in Python, a versatile and widely used programming language in the bioinformatics field. This choice ensures cross-platform compatibility and ease of integration with existing bioinformatics tools and libraries.

The CLI is designed to interact with a central shell script that orchestrates the execution of the pipeline. This script serves as the backbone of the FLYNC pipeline, managing the invocation of all child subprocesses and ensuring the sequential and conditional execution of various pipeline stages. The script provides visual cues from the pipeline, including progress updates for each main stage, allowing users to monitor the execution in real-time. It also redirects the output of the child subprocesses to a central logging file ‘run.log’. This approach enables users to maintain a record of the pipeline’s execution and facilitates debugging if necessary.

The CLI provides users with the ability to execute complex bioinformatics workflows through a series of simple, yet comprehensive commands. The FLYNC CLI offers multiple subcommands to accommodate various types of input data and user preferences: i) ‘run’: Executes the full pipeline using a YAML configuration file, allowing for a high degree of customization with minimal command-line arguments; ii) ‘srà: Facilitates the analysis of SRA accession numbers by running the full pipeline on a list of provided accession numbers; iii) ‘fastq’: Allows users to run the full pipeline on local FASTQ files, supporting both paired and unpaired read inputs. Each subcommand is equipped with a set of options to specify input files, output directories, metadata for differential expression analysis, and the number of computational threads to utilize. The FLYNC CLI is invoked using the ‘flync’ command followed by the desired subcommand and associated options. Examples of how to use the main subcommands: i) To run the pipeline with a configuration file: ‘flync run -c /path/to/config.yaml’; ii) To analyze a list of SRA accession number: ‘flync sra -l /path/to/list.txt -o /path/to/output/dir -m /path/to/metadata.csv -t 8’; iii) To process local FASTQ files: ‘flync fastq -f /path/to/fastq/dir -o /path/to/output/dir -p true -m /path/to/metadata.csv -t 8’.

The ‘-c/--config’ option is recommended as it simplifies the execution process by reading all necessary parameters from a YAML file. The ‘-l/--list’ option is mandatory for the ‘srà subcommand and requires a file with SRA accession numbers, one per line. The ‘-o/--output’ directory is advisable for all subcommands to specify where the results will be written. Metadata in CSV format is mandatory for differential expression analysis and is specified with the ‘-m/-- metadatà option. The number of computational threads can be adjusted using the ‘-t/--threads’ option to optimize performance based on available system resources.

The FLYNC CLI is designed with user-friendliness in mind, providing clear and concise help messages for each subcommand. Running ‘flync <subcommand> --help’ displays detailed information about the options available for that subcommand.

### FLYNC distribution and availability

FLYNC is distributed as a CLI tool to interact with the software pipeline and pre-trained ML model that classifies lncRNAs. To ensure accessibility and ease of use across different computing environments, FLYNC is available through multiple distribution methods.

FLYNC is available as a Docker image, providing a containerized environment that ensures consistency across different platforms. The Docker image can be pulled from the Docker Hub repository (hub.docker.com/r/rfcdsantos/flync). Users with ample storage space may opt for the “local-tracks” tagged image, which is larger in size but offers faster runtime due to the inclusion of UCSC tracks for feature extraction. When using Docker, it is recommended to map a local directory to the container to store results. This ensures that the output is accessible on the host machine after the Docker process is completed.

For users operating on Debian-based Linux systems, FLYNC can be installed locally using Anaconda. This method allows users to test, modify, and inspect the FLYNC scripts directly. The repository can be cloned, and the required environments can be set up using the provided ‘conda-env’ bash script.

FLYNC is distributed in a manner that prioritizes user convenience and reproducibility across different computing environments. The Docker and Conda distribution methods ensure that researchers can easily install and run FLYNC. FLYNC source code repository is hosted on GitHub (github.com/homemlab/flync) along with installation and usage instructions. Similarly, the SBUCELL module is available through GitHub (github.com/homemlab/subcell) with the same distribution approach.

### Bulk RNA-seq dataset

To assess the applicability of FLYNC, we utilized the publicly available bulk RNA-seq dataset GSE199164. This dataset encompasses brain gene expression profiles from *D. melanogaster* at three distinct post-eclosion ages: 3, 7, and 14 days. Notably, it includes data from both female and male subjects, providing a view of sexually dimorphic gene expression in fruit fly brains under normal conditions.

The dataset includes samples from different ages and sexes, providing a multifaceted biological context. This diversity allows us to demonstrate FLYNC’s adaptability in analyzing varying conditions within a single experiment. By applying FLYNC to this dataset, we can showcase the pipeline’s flexibility in handling complex experimental designs, including differential gene expression analysis across multiple biological variables. Additionally, this dataset also provided with a quick and reproduceable protocol to validate FLYNC findings through qPCR, with a short fruit fly growing step and easily collectable tissue.

### Single cell RNA-seq dataset

We also wanted to address the capacity of FLYNC to be applied to locally stored datasets. For that reason, we have resourced to scRNA-seq previously generated in our laboratory and already publicaly available(23). Also, considering the putative cell-type specific function of lncRNAs, this dataset provides sequencing data for an array of neural populations, including neural stem cells (neuroblasts) and differentiated neural cells (neurons). Hence, it would be possible to identify new putative lncRNAs, but also address its cellular context. Moreover, and as mentioned in the introduction, the brain, more than other organ already scrutinized, displays a tremendous amounts and diversity of lncRNAs, thus supporting the reasoning behind the selection of this dataset. Finally, validation of these results could be easily verified by sorting pure populations of neuroblasts or neurons and testing via RT-qPCR (see below).

### Fly husbandry

For experiments performed with whole heads, control, four 3-days old male flies (w1118) were collected. The whole head was extracted with forceps and immediately placed in ice-cold 250μL 1xPBS. For the experiments performed with sorted cells, labeling of neural stem cells and their progeny was done by using the stable line expressing both VT201094-Gal4 (VDRC; central brain and ventral nerve cord neuroblast driver) and UAS-myr::GFP. To maximize expression of GFP, fly husbandry was set up at 29°C. After approximately 5 days, third instar wandering larvae were collected for brain dissection.

### Brain dissociation and cell sorting

Approximately two hundred wandering third instar larvae were collected and dissected in supplemented Schneider’s medium (10% fetal bovine serum (Sigma), 20 mM Glutamine (Sigma), 0.04 mg/mL L-Glutathione (Sigma), and 0.02 mg/mL Insulin (Sigma), Schneider’s medium (Sigma)). Afterwards, brains were enzymatically dissociated in Schneider’s medium supplemented with 1mg/mL Papain (Sigma) and 1mg/mL Collagenase I (Sigma) for 1 hour at 30°C. Then, brains were washed twice with 1mL supplemented Schneider’s medium. To help pellet the brains between washes, samples were spun at 300 x g for 5 min. Subsequently, brains were resuspended in 200μL of supplemented Schneider’s medium and mechanically disrupted using a pipette tip. The cell suspension was filtered through a 30-μL mesh into a 5-mL FACS tube (BD Falcon) and immediately sorted by fluorescence activated cell sorting (FACS) (FACS Aria II, BD). GFP-positive cells were collected in 750μL of supplemented Schneider’s medium. Since NBs and their progeny can be distinguished by their size, NBs and neurons were collected separately. Once cells were collected, the volume of the tube was corrected to a final volume of 1000μL. Cells were then immediately used for RNA extraction.

### RNA extraction and qPCR

mRNA was isolated using TRIzol™ LS Reagent (Invitrogen) according to the manufacturer’s instructions. RNA was then treated with TURBO DNA-free™ Kit (Invitrogen™) to ensure removal of remnant DNA. cDNA was prepared using the RevertAid First Strand cDNA Synthesis Kit (Thermo Scientific™). qPCRs were done using GoTaq qPCR Master mix (Promega) on a QuantStudio™ 5 Real-Time PCR System (Applied Biosystems™). Expression of all genes was normalized to act5C or vglut and relative levels were calculated versus control using the ΔCt method(24). All measurements were done with technical triplicates. A list of the primers used can be found in Table 1.

**Table 1–.**
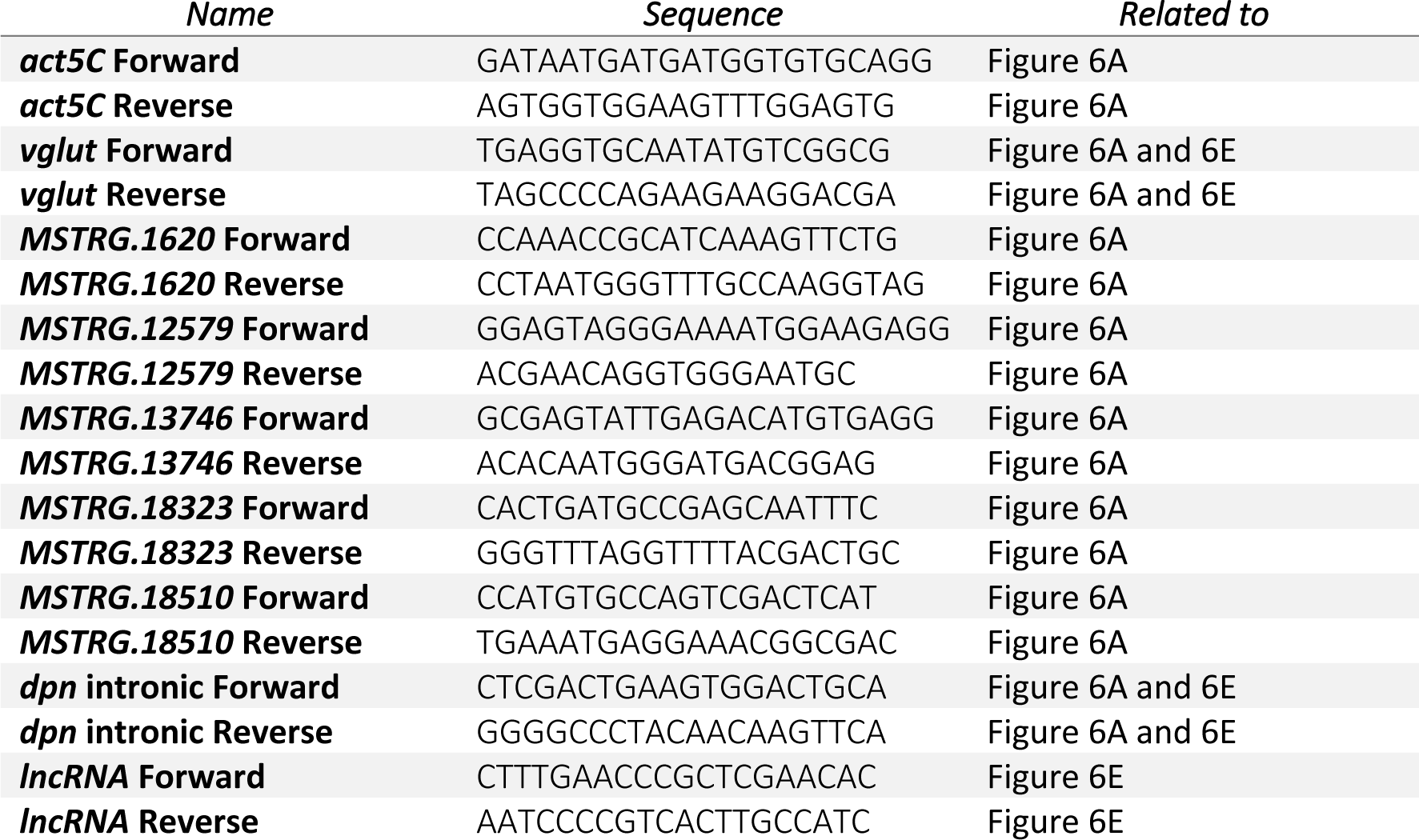
List of primers used for RT-qPCR experiments.

## RESULTS AND DISCUSSION

### The *Drosophila melanogaster* non-coding genome annotation gap

Genome-wide sequencing approaches have been extensively applied to virtually all model organisms across many scientific disciplines. *Drosophila melanogaster* is one of the most used animal models, including in the study of important human diseases such as Alzheimer’s and brain tumorigenesis(25).

Non-coding RNAs have increasingly been implicated in critical molecular mechanisms of disease, but *D. melanogaster* non-coding genome has been understudied (26). We compared the Protein Coding Gene (PCG) and Non-coding Gene (NCG) counts in genome annotations of several model organisms used in research (Figure 1A). It is noticeable that organisms widely used in RNA biology studies exhibit a higher percentage of annotated NCG. In *Caenorhabditis elegans*, where the RNAi was discovered and characterized, NCGs account for more than half (55%) of the total genes annotated to date. In contrast, annotated NCGs *D. melanogaster* show only a fraction of this (22%) (27). This argues in favor of a multitude of non-coding RNAs yet to be discovered in *D. melanogaster* potentially hiding a layer of RNA regulatory circuits.

**Figure 1–.**
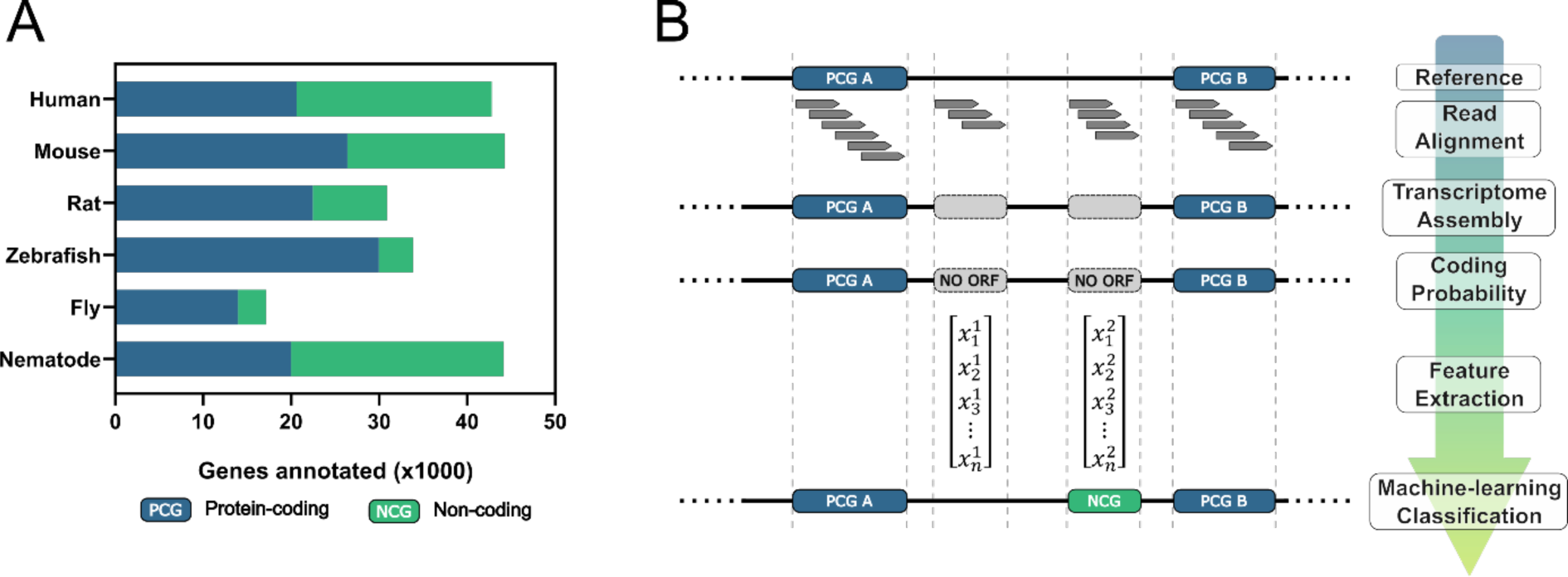
Bridging the annotation gap of non-coding RNAs in D. *melanogaster* with FLYNC. (A) Comparative analysis of Protein Coding Gene (PCG) and Non-coding Gene (NCG) counts in genome annotations across various model organisms. This figure illustrates the lower percentage of annotated non-coding genes in *Drosophila melanogaster* compared to other species. (B) Schematic representation of the FLYNC machine learning approach for classifying RNA transcripts. The process includes building gene models from sequencing reads, evaluating non-coding potential, constructing a feature matrix, and inferring classifications using a pre-trained ML model.

The scarce bioinformatics toolset to study NCGs in *D. melanogaster* has led us to the development of FLYNC. FLYNC is our effort to close the gap between the non-coding and coding *D. melanogaster* genome annotation, through an evidence-based gene discovery and ML-driven classification approach. The high-level overview of FLYNC approach is to: i) build gene models from transcriptomic sequencing reads; ii) evaluate the non-coding potential of their transcripts; iii) construct an evidence-based feature matrix for each non-coding gene model; iv) inference a pre-trained Machine Learning (ML) model with each gene and classify them as a previously unannotated RNA gene (Figure 1B). This approach feeds the ML model experimentally supported targets, which ultimately enriches the pool of true positive hits and helps validate candidate genes.

### Overview of FLYNC

FLYNC was implemented as an end-to-end pipeline to facilitate its usage and abstract some technical requirements away from the end users. FLYNC can be conceptually segregated into two main stages, a Bioinformatics stage (BI) and an Artificial Intelligence inference stage (AI) (Figure 2). The BI stage implements a bioinformatics pipeline based on the Tuxedo2 protocol (28) coupled with a coding potential assessment using CPAT (29). The AI stage queries the pre-trained ML model to refine the outputs of the BI stage by classifying them as new lncRNAs.

**Figure 2–.**
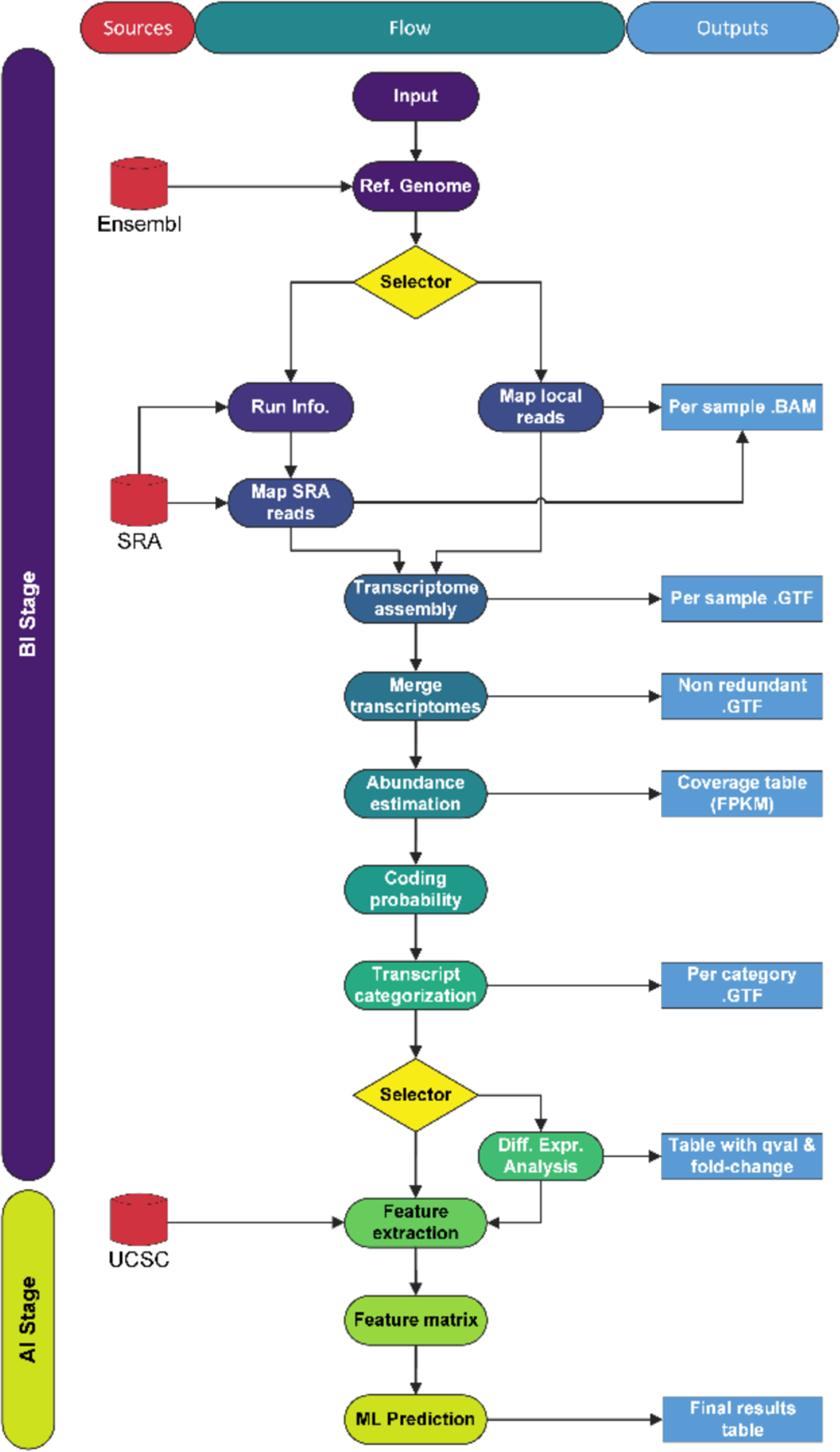
Overview of the FLYNC bioinformatics pipeline. The two main stages are depicted: Bioinformatics (BI) and Artificial Intelligence (AI). The figure details the data flow from input through various analytical steps to the final output, highlighting the integration of external data sources (left) and the pipeline’s output components (right).

The program depends on critical information from three different external public sources (Figure 2 – under “Sources”): Ensembl for the reference genome sequence and annotation; Sequence Read Archive for retrieving published transcriptomic reads; and the University of California Santa Cruz: Genomics Institute hosting the genome browser tracks (BigBed and BigWig files) used to build the features for the predictive model. During the computational steps, various files are created that may be used in future runs or fed into other bioinformatic pipelines (Figure 2 – under “Outputs”). The final results file comprises vital information to explore, filter, analyze and act on the recently discovered candidate lncRNA genes.

The program will parse the user-given instructions (**INPUT**) to start the pipeline with the desired configuration. This requires: i) the sequencing reads that will be mapped to the reference genome. These can either be “fastq” files supplied by the user or SRA accession numbers that will automatically download and temporarily store the reads locally; ii) the path to store the results of the pipeline. Optionally, a sample metadata file can be supplied to map each sequencing sample to a condition (e.g. wt *vs.* mutant). This is used to run a differential gene expression analysis (DGE) on the newly assembled transcripts (Figure 2 – under “Flow”). Using this configuration, the program starts the pipeline by downloading the reference genome annotation and sequences to use at runtime (**REF. GENOME**). Based on the initial configuration, the program will either (**SELECTOR**): i) map locally stored sequencing reads using the “fastq” subcommand (**MAP LOCAL READS**); ii) map remotely stored sequencing reads retrieved from the SRA database along with vital SRA run metadata using the “sra” subcommand (**RUN INFO.** & **MAP SRA READS**). After all samples are mapped, the results are used to create an annotation of each sample’s transcriptome (**TRANSCRIPTOME ASSEMBLY**) which is then merged into a single non-redundant transcriptome annotation of all selected samples (**MERGE TRANSCRIPTOMES**). The quantity of each transcript is estimated (**ABUNDANCE ESTIMATION**) followed by an assessment of ORF presence (**CODING PROBABILITY**). The binarization of the previous step – “coding” vs “non-coding” – combined with the possible gene models of the new (previously unannotated) transcripts allowed us to develop a script that categorizes these candidate genes (**TRASCRIPT CATEGORIZATION**). Namely, into different subclasses of both coding (new gene isoforms, new micro ORFs) and non-coding (new lincRNAs, intronic lncRNAs, antisense lncRNAs). To finalize the BI stage, the optional step (**SELECTOR**) of running a Differential Expression Analysis (**DIFF. EXPR. ANALYSIS**) on the samples may be used to help refine the final output of the program. For each new candidate lncRNA transcript a set of features is pulled from the UCSC database (**FEATURE EXTRACTION**) and combined into a unified feature matrix of feature values per candidate lncRNA transcript (**FEATURE MATRIX**). Finally, we feed this matrix to our predictive model and store the result into a final dataset (**ML PREDICTION**).

### Training dataset and feature engineering

lncRNAs are typically described as transcript with over 200 nucleotides that is not translated. However, the observation of the absence of an ORF in a genome-wide scale is often insufficient to provide strong leads for further experimental characterization (30). We reasoned that transcriptomic studies focused on different genomic properties – such as transcription factor binding site distribution or sequence conservation – could be leveraged by AI algorithms to identify hidden patterns and help discover novel long non-coding transcripts. UCSC provides a curated Tracks Hubs with multiple high-quality tracks available for the *Drosophila melanogaster* genome. We looked at these data for possible features to train an AI model and included relevant UCSC tracks in our pre-training dataset. This allowed us to: ii) engineer features and estimate correlations; ii) refine feature selection for the final training dataset; iii) evaluate the most suitable AI algorithm for the objective.

The final features used to train the AI model are shown in Figure 3A depicting an example of a genomic region to visually represent the source track data. The suitable metric extracted from each feature is presented on the right side. The metric was chosen as a representation of what each value might represent biologically for a given pair of coordinates (boundaries of a gene). For example, the “modENCODE CAGE samples” (bottom track) maps Transcription Start Site (TSS) peaks. Therefore, only the maximum value for each coordinate pair was considered, which can be interpreted as the presence and strength of a TSS. Similarly, the “GC percentage in 5-base windows” (top track) represents the GC% that correlates with DNA stability and thus, the mean GC% content of each coordinate pair was extracted.

**Figure 3–.**
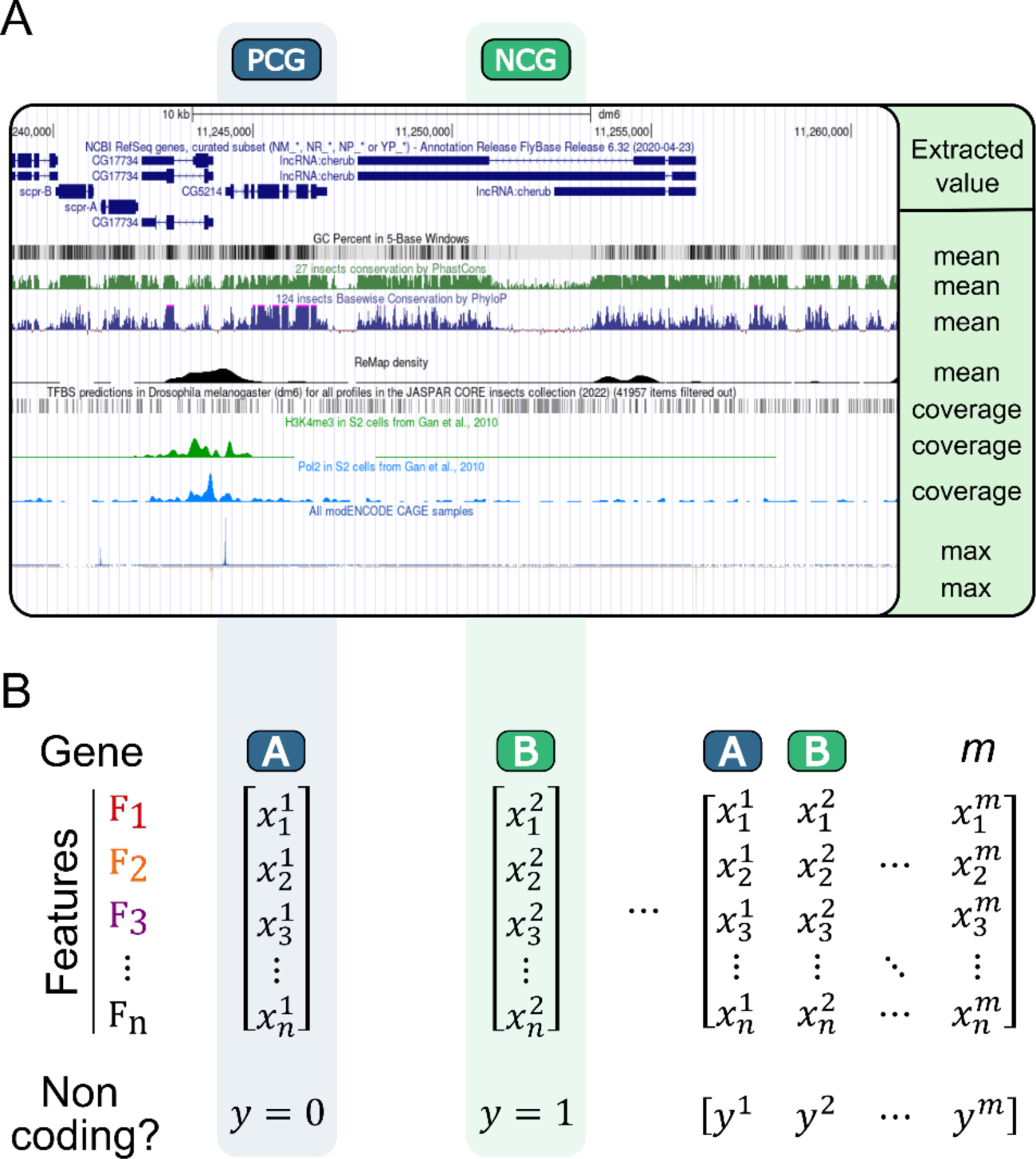
Building a gene feature matrix for FLYNC ML. Visualization of the genomic features used to train the FLYNC ML model, with an example genomic region displayed. Each track represents a different feature, such as transcription start sites or GC content, with the extracted metric for each feature described on the right side. The metrics of each feature were extracted for each gene and the target label assigned: 0 = Protein Coding Gene (PCG); 1 = Non-Coding Gene (NCG).

Our biologically driven feature engineering approach resulted in a total of 9 features used to train the ML model: i) sequence length; ii) mean phyloP conservation score (124 insects); iii) mean PhastCons score (27 insects); iv) maximum modENCONDE CAGE samples (100 bp flanking region); v) maximum modENCONDE CAGE samples (gene span); vi) coverage of JASPAR CORE; vii) mean of ReMap score; viii) mean GC percent; ix) coverage of RNA Polymerase 2 (31–36). A schematic representation of the feature matrix used to build the training dataset is shown in Figure 3B.

To prepare the final training dataset a .BED file of all *D. melanogaster* annotated genes (BFGP6.32 v103) was prepared by filtering out all tRNAs, rRNAs and sRNAs genes which excludes non-lncRNAs. This file was later used to extract the above features from the source database (UCSC) and compile them into a tabular dataset. To assign the target variable (label) for the machine learning model, all genes classified as “ncRNA” were assigned a value of True, while all others were assigned a value of False. The clear binary classification nature of the problem has the aim of accurately predicting whether a given transcript is a long non-coding RNA (lncRNA) or not. The complete training dataset is available in the supplementary materials. (Supp. Materials).

In selecting an appropriate AI programming methodology, we prioritized those that offered an optimal balance between performance and explainability. Consequently, we excluded unsupervised AI methods, which are often perceived as “black boxes”, and instead focused on supervised and explainable machine learning frameworks (37). To quickly test multiple AI algorithms we prepared a pre-training dataset comprising a small, balanced sample of 10,000 randomly selected *D. melanogaster* NCG and PCG coordinates, from which we extracted the corresponding UCSC track values. Using the ‘lazypredict’ python library we conducted five independent training sessions for each model, reserving 25% of the dataset per session to calculate each model’s accuracy. Results from the 5 most accurate algorithms is provided in the supplementary materials (Supp. Materials). The top-performing machine learning algorithm was the Random Forest (RF), with an accuracy of 0.8651±0.0049, which we subsequently adopted for further data science tasks.

### Model training and evaluation

The Random Forest algorithm was selected as the optimal approach for achieving superior prediction scores and providing greater transparency into the model’s prediction of the feature set outcomes. RF models are a form of ensemble learning method based on the outcome of multiple Decision Trees (DT) in which each DT is constructed from a random sample drawn from the training data (38). During the prediction phase, the RF model outputs the class that is agreed upon by the majority of the individual trees. RF models are particularly adept at handling large datasets with high dimensionality and dealing with missing or unbalanced data. They are also suited to handle a variety of data types, including numerical, categorical, and mixed data types, and are especially robust for managing large datasets with numerous features (39). There are several advantages to using Random Forest models, including i) relative ease of use and interpretation, making them accessible to non-experts; ii) robustness to noise, outliers, and missing or unbalanced data; iii) quicker training times and less resource intensive. Combined, these RF features are uniquely fitted to the overall purpose of this work: to establish an AI-powered bioinformatics framework to streamline *D. melanogaster* lncRNA discovery and study. Consequently, the community will be able to quickly retrain the model to add new features or when new lncRNA are annotated.

The Random Forest (RF) model was trained using the complete *D. melanogaster* genome. It is pertinent to note that the dataset employed in this training exhibited a significant class imbalance, with a notably higher proportion of non-long noncoding RNA (not-lncRNA) instances compared to long noncoding RNA (lncRNA) 228612 and 7665, respectively. Given the extensive and intricate array of biologically characteristics that distinguishing the two classes, we posited that preserving this imbalance would be advantageous for the model’s performance. Specifically, that the imbalance would skew the model towards achieving greater Precision as opposed to Recall. This strategic decision was anticipated to result in a diminished yield of positive predictions from the model. Such an outcome is serving a dual purpose: it narrows the field of potential study subjects and concurrently bolsters the reliability of the selected candidates for subsequent experimental validation.

To assess the performance of the RF model, we used the 5-fold cross validation technique, which splits the data into five equal subsets and iteratively trains the model on four subsets and tests it on the remaining one. We computed the accuracy, precision, recall, and F1-score metrics for each fold using the whole training set and averaged them over the five folds. The results are shown in Figure 4A, with the standard deviations indicated by the error bars. The RF model achieved a very high accuracy of 0.9847±0.0004, indicating that it correctly classified most of the instances. The precision score was 0.9279±0.0079, meaning that the model had a low rate of false positives. The recall score was 0.5721±0.0142, meaning that the model captured more than half of the true positives. The F1-score, which is the harmonic mean of precision and recall, was 0.7077±0.0118, indicating a good balance between the two metrics.

**Figure 4–.**
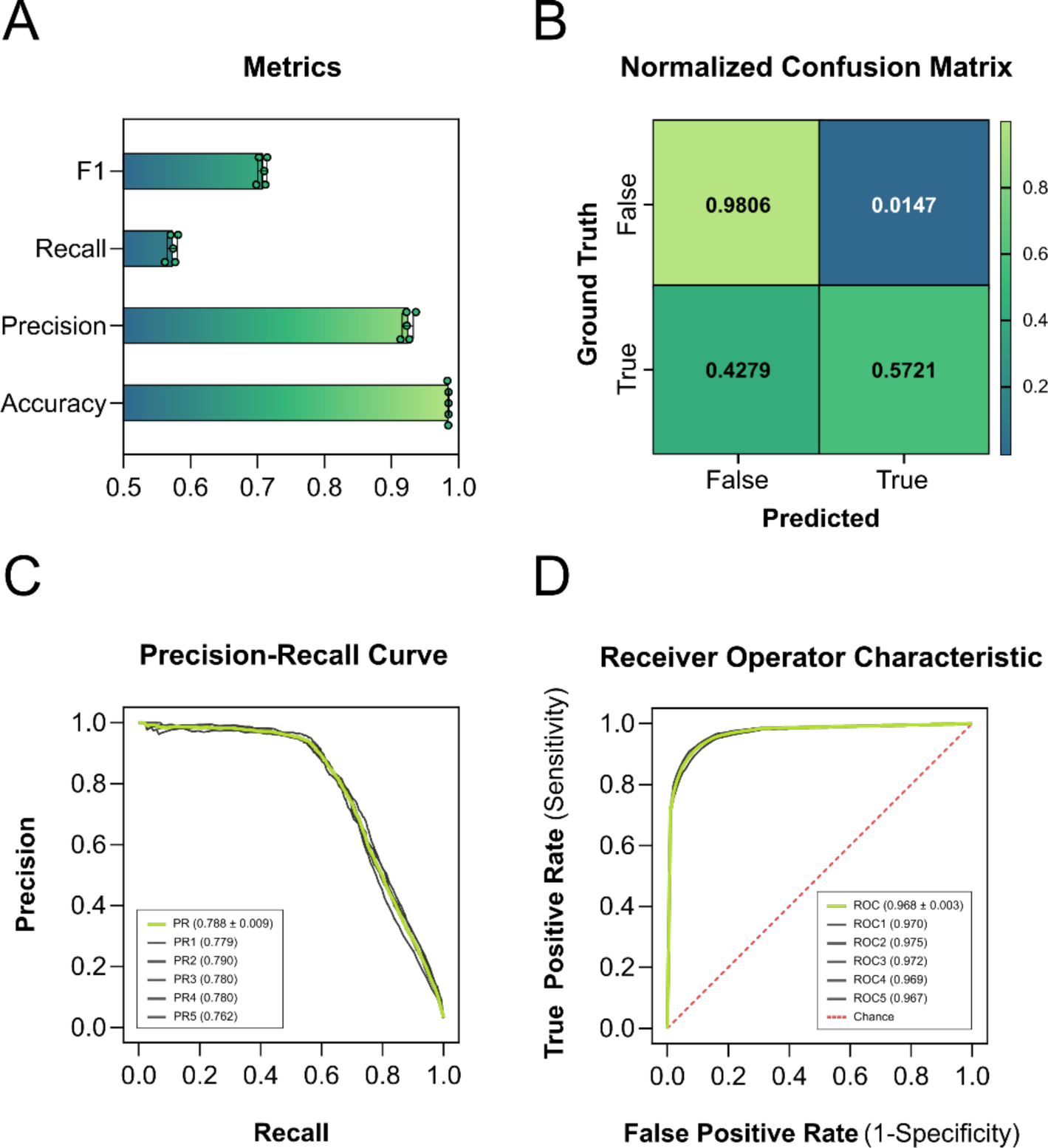
Evaluation of FLYNC Random Forest model across 5-fold cross-validation metrics. (A) Graphical representation of the Random Forest (RF) model’s performance across four common metrics: F1-score, Recall, Precision and Accuracy with 0.7077±0.0118, 0.5721±0.0142, 0.9279±0.0079 and 0.9847±0.0004, respectively. (B) Normalized confusion matrix for the RF model, showing the proportion of true and false predictions for non-lncRNA and lncRNA instances, emphasizing the model’s precision in classification. (C) Precision-recall curve for the Random Forest model, depicting the trade-off between precision and recall across different thresholds with area under the curve (AUC) value of 0.788±0.009. (D) Receiver Operating Characteristic (ROC) curve for the Random Forest model, displaying the relationship between the true positive rate (TPR) and false positive rate (FPR) at various thresholds, with an AUC of 0.968±0.003.

The confusion matrix, which shows the distribution of the predicted labels versus the true labels, was normalized by dividing each cell by the total number of instances in each class. This allows us to compare the performance of the model across different classes, regardless of their size. The normalized confusion matrix is shown in Figure 4B, indicates that the model rarely misclassified non-lncRNA as lncRNA or vice versa. Namely, that it almost perfectly identified all the non-lncRNA instances and detected more than half of the lncRNA instances.

To further evaluate the performance of the RF model, we plotted the precision-recall curve and the receiver operating characteristic (ROC) curve, which are two common ways of visualizing the trade-off between true positives and false positives. The precision-recall curve shows how the precision and recall vary as the threshold for positive prediction changes. The area under the curve (AUC) is a measure of how well the model performs across all possible thresholds. The RF model achieved an AUC of 0.788±0.009, which is relatively high given the class imbalance in the data. The ROC curve shows how the true positive rate (TPR) and the false positive rate (FPR) vary as the threshold for positive prediction changes. The AUC for the ROC curve is also a measure of how well the model performs across all possible thresholds. The RF model achieved an AUC of 0.968±0.003, which is very close to the ideal value. The precision-recall curve and the ROC curve are shown in Figure 4C and 4D, respectively.

The RF model performance metrics are comparable with those of widely used ML-based lncRNA identification models (29, 40, 41), and other lncRNA-related models (42, 43). Even though, such metrics are often not easily comparable since some are calculated with a random balanced subsample of the training data or were tested on different species. Unfortunately, very few of these models are usable on *D. melanogaster* data, with the vast majority focusing on the Human and Mouse organisms (44). Moreover, the tools that do provide species-agnostic capabilities often require high technical knowledge and/or a model retraining step. Instead of trying to outperform previous and valid methods, our approach leverages CPAT, one of the few tools pre-trained on *D. melanogaster* data with non-overlapping FLYNC features or methods. CPAT employs a Logistic Regression (LR) algorithm on RNA sequence features (ORF, Fickett score and Hexamer) while FLYNC’s ML classifier employs an RF algorithm mainly on experimental data (43). Using CPAT to filter out protein coding transcripts, FLYNC’s classifier then refines the remaining subset based on nucleotide distribution (GC content), sequence size and conservation (PhastCons & PhyloP), transcriptomic signals (CAGE & Pol2) and regulatory context (ReMap & JASPAR) patterns of each transcript.

Overall, these results indicate that the RF model is highly reliable in distinguishing non-lncRNA from lncRNA, and that it can effectively narrow down the candidates for lncRNA discovery and validation.

### Applying FLYNC to publicly available bulk transcriptomic data

We applied FLYNC to a publicly available bulk transcriptome sequencing dataset (GEO accession GSE199164) and investigated the lncRNA expression patterns in different groups of *D. melanogaster* samples. This accession contains sequencing data from 3, 7, and 14-day old male and female flies. We selected six samples of 3-day old Oregon wild-type flies, with three male and three female replicates and provided their metadata to FLYNC for differential gene expression (DGE) analysis (Table 2). We used the command line interface (CLI) of FLYNC to run the workflow on these samples, as described in detail in the Materials and Methods section. The command was: ‘flync sra --list list.txt --metadata metadata.csv --threads 8 --output bulk_run’. This command instructed FLYNC to use the ‘srà subcommand for SRA accessions, to read the sample list and metadata from the files ‘list.txt’ and ‘metadata.csv’, respectively, to use eight CPU threads for parallel processing, and to store the results in a folder named “bulk_run”. The result for this run is provided as supplementary material files.

**Table 2–.** Phenotypic metadata for each SRA accession used in bulk RNA-seq applicability experiments. The table presents the metadata for two experimental setups: ‘Male *vs.* Female’ and ‘3-day *vs.* 7-day,’ reflecting the diverse conditions tested using the same transcriptomic dataset.

| *Male vs Female* |  | *3-day vs 7-day* |  |
| --- | --- | --- | --- |
| **SRR18432803** | *Male* | **SRR18432803** | *3-day* |
| **SRR18432804** | *Male* | **SRR18432804** | *3-day* |
| **SRR18432805** | *Male* | **SRR18432805** | *3-day* |
| **SRR18432812** | *Female* | **SRR18432812** | *3-day* |
| **SRR18432813** | *Female* | **SRR18432813** | *3-day* |
| **SRR18432814** | *Female* | **SRR18432814** | *3-day* |
|  |  | **SRR18432800** | *7-day* |
|  |  | **SRR18432801** | *7-day* |
|  |  | **SRR18432802** | *7-day* |
|  |  | **SRR18432809** | *7-day* |
|  |  | **SRR18432810** | *7-day* |
|  |  | **SRR18432811** | *7-day* |

FLYNC identified a total of 20763 assembled transcripts that were not previously annotated. After applying the CPAT step, 9990 transcripts were identified as possible non-coding transcripts. FLYNC ML classified 1141 of such transcripts as lncRNAs with a probability greater than 50%, of which 167 with a probability greater than 75%.

One of the advantages of FLYNC is its flexibility and adaptability to different experimental settings. For instance, we performed a different analysis using the same GEO dataset by grouping the samples by age instead of sex. Specifically, we compared the lncRNA expression profiles of 3-day old and 7-day old flies, both male and female, to identify age-related lncRNA differences. We ran the FLYNC pipeline with the same command as before but updated the accessions in list.txt and changed the metadata.csv file to reflect the new grouping criteria (Table 2).

In this second call, FLYNC identified a total of 27,021 previously unannotated transcripts. Of those, 11929 were identified as non-coding by CPAT. FLYNC ML classified 886 as lncRNAs with a probability greater than 50%, and 118 with a probability greater than 75%.

We have shown how FLYNC can be easily applied to different biological questions and generate novel insights into lncRNA function and regulation. FLYNC is a powerful tool that offers flexibility and adaptability to different experimental settings and can easily incorporate data from different public transcriptomic datasets. For instance, other datasets with 3-day old flies could be used to expand our analysis, and it would only require the user to add the required accessions and reflect the phenotypic data in the metadata.csv file. FLYNC is also able to reduce the number of candidate lncRNA genes for downstream studies. Notably, FLYNC ML classfier led to a reduction of 88,6% (1141/9990) and 92,6% (886/11929) in candidate lncRNAs (>50% probability) for the first and second call, respectively. This will help streamline the discovery and characterization of novel lncRNAs in *D. melanogaster*. In conclusion, FLYNC is a valuable resource for the lncRNA research community and can facilitate the identification and analysis of lncRNAs in different conditions and contexts.

### Applying FLYNC to single-cell transcriptomic data

We aimed to expand the applicability of FLYNC to single-cell RNA sequencing (scRNA-seq), one of the latest and most powerful sequencing experiments. Importantly, this allows to withdraw cell-type specific information, something not possible from highly complex and heterogenous samples, as the example above. Typically, the output of the last step of the scRNA-seq pipeline is a matrix of gene expression values for each cell. This matrix can be used to identify different cell types and states, as well as to study gene expression patterns and regulatory networks (45). The Seurat R package is one of the most popular and widely used scRNA-seq analysis tools (46). However, the results are tied to the original annotated reference genome (for example, when using the 10X Chromium platform and the Cell Ranger software). Furthermore, we faced a challenge when trying to identify raw reads of each individual cell with the goal of using FLYNC. Since these reads are not easily accessible from, we developed a performant program named “SUBCELL” (https://github.com/homemlab/subcell).

SUBCELL takes a folder with the original FASTQ files of the scRNA-seq experiment, and the cell barcodes extracted with the ‘WhichCells()’ method of the Seurat R package as arguments. It then scans through each FASTQ file trying to find the specific cell barcode in the FASTQ header, and groups each matching read to a new FASTQ file, effectively grouping reads of the same cluster, as analyzed with Seurat. These reads could now be used on other bioinformatics pipelines that require a new alignment step or de novo assembly, such as FLYNC (Figure 5A).

**Figure 5–.**
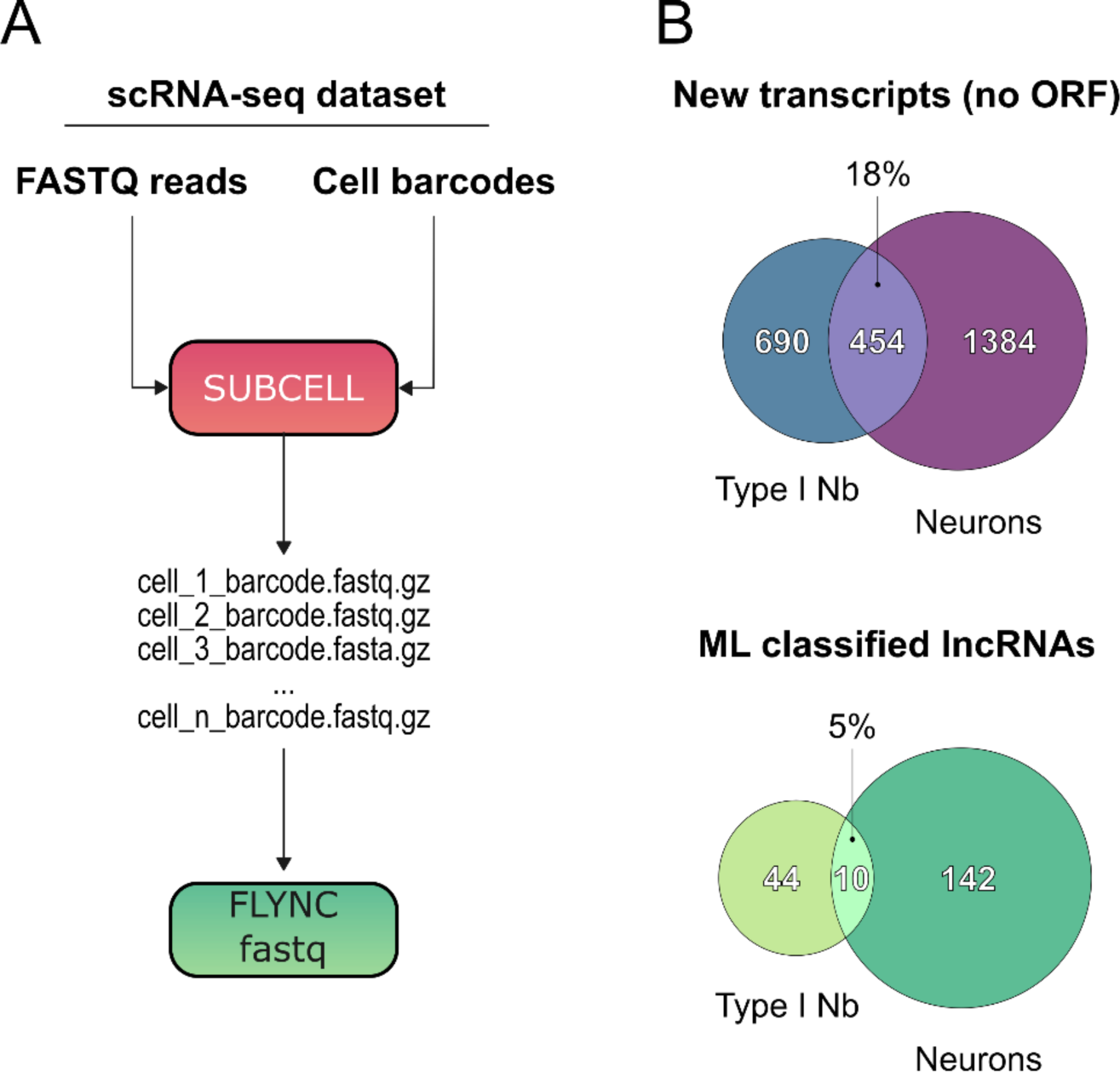
Applying FLYNC to single cell RNA-seq data with SUBCELL pre-processing step. (A) Diagram of the SUBCELL program’s functionality, which groups sequencing reads by cell barcode from the original FASTQ files, enabling their use in FLYNC ML pipeline for further analysis. (B) Summary of the FLYNC pipeline application to single-cell RNA-seq data for Type I Neuroblasts and Neurons, showing the number of identified non-coding transcripts and the reduction in candidate lncRNAs achieved by the ML classifier.

To demonstrate this, we extracted the cell barcodes for Type I Neuroblasts (Nb) (*D. melanogaster* brain stem cells) and the barcodes of Neurons (*D. melanogaster* differentiated brain cell type) using the ‘WhichCells()’ method into two files: ‘neuroblasts.barcodes’ and ‘neurons.barcodes’, respectively. We ran SUBCELL with these files to group together reads from each cell barcode and output those into cell type-specific folders of FASTQ reads. We then ran the FLYNC ML pipeline on these two sets of transcriptomic reads. The individual cell reads were then given by the FLYNC ML pipeline using the ‘fastq’ subcommand.

A diagram summarizing both FLYNC calls for Type I Nb and Neurons is shown in Figure 5B. The total number of possible new non-coding transcripts as output by the CPAT step is 1,144 for Nb and 1,838 for Neurons, with 454 (18%) overlapped in both analyses. FLYNC ML classifier identified 54 of 1,144 as lncRNAs in Nb and 152 of 1,838 as lncRNAs in Neurons, with an overlap of only 10 (5%). This effectively reduced the universe of candidate lncRNAs in 95,3% and 91,7% for Nb and Neurons, respectively. Additionally, this comparative analysis shows that the non-overlapping candidate lncRNAs could represent cell-type specific lncRNAs involved in maintaining an explicit cell state, as undifferentiated Nb or fully differentiated Neurons.

FLYNC maintained its ability to reduce the universe of lncRNAs as shown when applied to bulk transcriptomic studies with overall values (bulk and scRNA-seq) values ranging 88-95% of CPAT identified non-coding transcripts. Notably, this reduction is accomplished by the machine learning classifier based on features extracted from genome-wide experiments. Hence increasing the confidence of positive hits and providing a platform for successful molecular characterization experiments.

### Validation of FLYNC-identified putative lncRNAs

In the context of our analysis pipeline, wherein both bulk and scRNA-seq datasets were investigated, a meticulous validation of the putative lncRNAs identified was performed. This validation process involved specific experiments tailored to the characteristics of the two distinct datasets used. First, whole RNA extraction from 3-days old adult male heads, followed by DNAse treatment, was carried and used to characterize the occurrence of lncRNAs disclosed by FLYNC (Figure 6A and Table 1). Putative lncRNAs tested were selected based on (i) their potential as lncRNAs; (ii) their level of expression within the published dataset; (iii) and their relative genomic proximity to neural-related genes. Expression of *act5C* and *vglut* genes served as positive controls, and results were normalized to the expression of the *vglut* gene. Intronic *dpn* primers were used to completely exclude potential DNA contamination (negative control). Furthermore, a criterion was applied for positivity, defining expression as amplification occurring before cycle 35; any amplification beyond this threshold was considered non-expressed/non-detectable. As depicted in Figure 6A, three out of the five putative lncRNAs subjected to analysis (MSTRG.1620, MSTRG.12579, and MSTRG.13746) demonstrated expression (C_t_ < 35), thereby corroborating their identification by FLYNC. In contrast, lncRNAs MSTRG.18323 and MSTRG.18510 exhibited amplification beyond the defined threshold, despite amplifying much earlier than the negative control (*dpn* intronic). This is indicative of either lack of transcription or expression levels below the limits of detection. Importantly, several lncRNAs are notorious for their low expression(18). For that reason, the impossibility to identify them by RTqPCR does not invalidate their existence or biological relevance, namely in a complex sample where each individual lncRNA might be underrepresented.

**Figure 6–.**
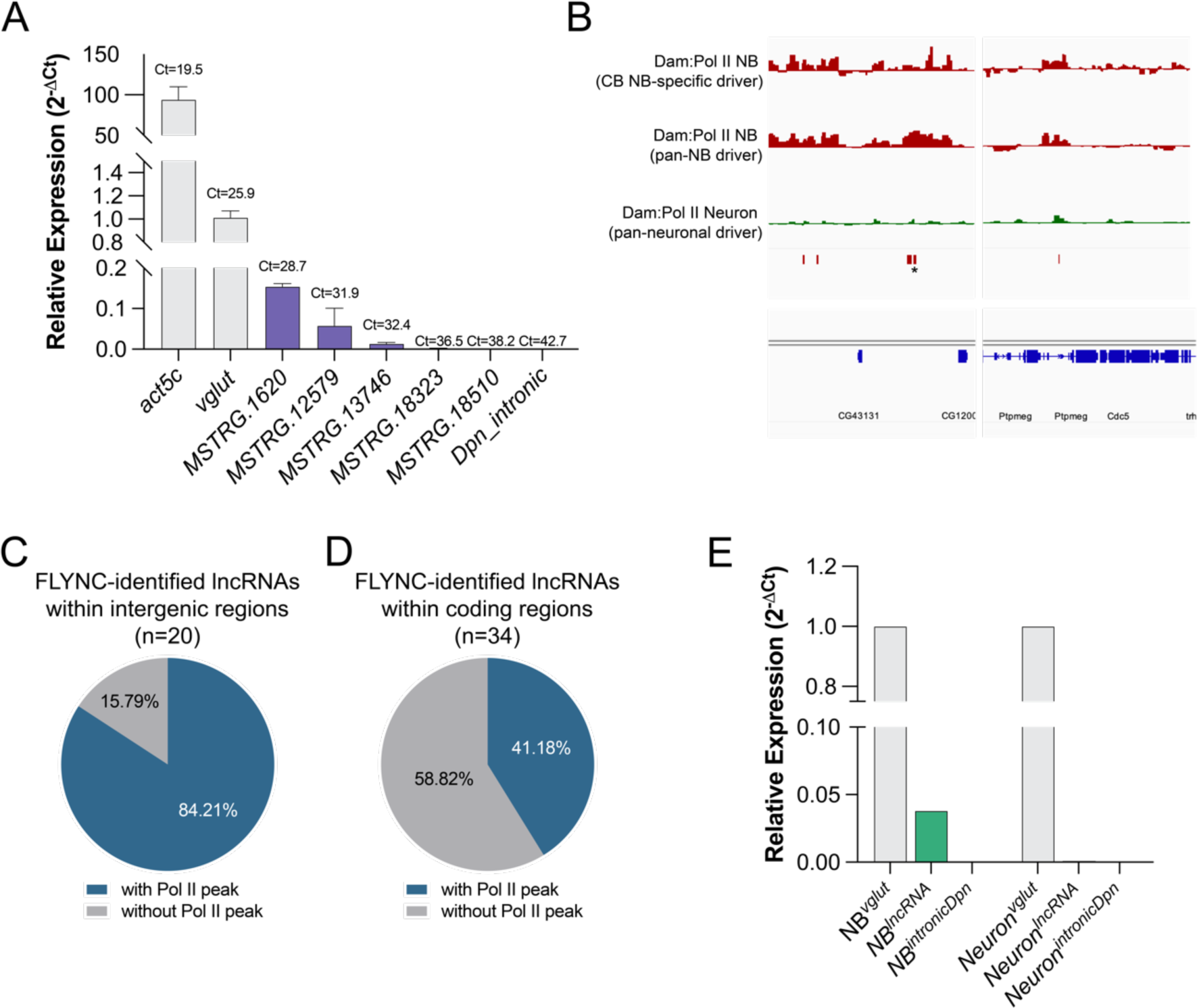
*In vivo* validation of FLYNC-identified lncRNAs. (A) RTqPCR experiments performed in RNA extracted from 3-days old adult male whole heads. Tested putative lncRNAs were select based on the probability of being a lncRNA and their expression levels. *act5C* and *vglut* expression was used as positive controls. Intronic *dpn* primers were used as negative control. Results were normalized to the expression of *vglut*. Experiments were performed with 3 biological replicates. (B) Representative genome browser tracks for targeted DamID experiments to evaluate RNA Pol II recruitment in neuroblasts (red tracks) and neurons (green tracks). Genomic location of neuroblast-specific putative lncRNAs identified by FLYNC are indicated with red boxes. (C-D) Analysis of RNA Pol II peaks of putative lncRNAs located within (C) intergenic and (D) coding regions for all identified putative lncRNAs. (E) RTqPCR experiments performed in RNA extracted from purified populations of either neuroblasts (NB) or neurons to evaluate the expression of the putative NB-specific lncRNA identified by FLYNC. *vglut* expression was used as positive control. Intronic *dpn* primers were used as negative control. Putative lncRNA tested in indicated in panel B with an asterisk(*).

In parallel, we have tested whether the putative lncRNAs identified using the scRNAseq dataset would also have some biological significance. To streamline this analysis, particular emphasis was directed toward lncRNAs exhibiting NB-specific expression. The reasoning behind this approach is two-fold: validate the existence of the lncRNA by an independent method and corroborate its cell-type specificity. Initially, we scrutinized the recruitment of RNA Polymerase II (Pol II), as determined by Targeted DamID, to the genomic *loci* of these individual lncRNAs (Figure 6B). The genomic locations of these lncRNAs are demarcated by red vertical boxes. As observed, some of the putative lncRNAs show Pol II recruitment with high specificity (peaks observed in NBs – red genome browser tracks –, while conspicuously absent from neurons – green genome browser tracks) (Figure 6B, left panel). However, in several instances, these Pol II peaks are either absent or non-specific, being prevalent in both NB and neurons (Figure 6B, right panel). In fact, when all 54 putative lncRNAs are analysed, it is possible to observe two distinct patterns. On one hand, we have observed that putative lncRNAs identified with intergenic regions correlate better with Pol II recruitment, with approximately 84% presenting Pol II peaks in a cell-type specific fashion (Figure 6C). On the other hand, FLYNC-identified lncRNAs located within coding regions present low correlation and specificity with Pol II recruitment (Figure 6D). Likely, this is because we do not have information about directionality of the identified RNAs, considering that scRNA-seq dataset used is not strand-specific. Hence, Pol II binding probably reflects the expression of the coding gene, and not necessarily a lncRNA. That, in addition to low or absent expression, can account for the lack of Pol II peaks in those regions. Nevertheless, to validate the presence of NB-specific lncRNAs identified within the scRNA-seq dataset, we have decided to select one located within an intergenic region and that showed NB-specific Pol II peaks (Fig. 6B, labelled with an asterisk). To test the expression of the putative lncRNA, we used the central brain and ventral nerve cord NB-specific Vienna Tile Gal4 line #201094 to drive the expression of CD8::GFP specifically in NBs. Since GFP presents significant stability, it is inherited by the NB progeny, including neurons, efficiently labelling them. Approximately 200 wandering third instar larvae brains were dissected, dissociated, and cells sorted by FACS, based on their fluorescence and size, to clearly distinguish NBs from neurons. NBs and neurons were collected separately, and RNA was extracted. As mentioned before, to avoid any DNA contamination that could hinder our analysis, samples were treated with DNase. Additionally, *vglut* expression was used as a positive control and *dpn* intronic primers used as negative control. Importantly, we were able to detect levels of the transcript and, as predicted by FLYNC, this lncRNA is NB-specific, with neuronal expression below the detection limits (Figure 6E).

## CONCLUSION

In this study, we have addressed a significant gap in the annotation of the non-coding genome of *D. melanogaster*, a model organism paramount to the understanding of highly conserved developmental pathways and human disease mechanisms. Despite its widespread use, the *D. melanogaster* non-coding genome has remained underexplored, particularly in comparison to other model organisms. Our findings underscore the potential for a vast array of non-coding RNAs yet to be discovered, which may play crucial roles in RNA regulatory networks. To bridge this gap, we developed FLYNC, an innovative bioinformatics pipeline that leverages a machine learning-driven classification approach to identify and annotate non-coding genes. FLYNC integrates a robust bioinformatics stage, based on established protocols and coding potential assessment, with an AI inference stage that refines and classifies the outputs as new lncRNAs. The pipeline effectively utilizes data from Ensembl, the Sequence Read Archive, and UCSC Genomics Institute, ensuring a comprehensive and evidence-based approach to gene discovery.

The Random Forest model, central to the AI stage of FLYNC, showed exceptional performance with high accuracy and precision. Its performance is comparable to other existing lncRNA identification models, with the benefit of being able to handle class imbalances, noise and outliers. The model’s transparency and interpretability will facilitate its adoption by the research community, and hopefully provide a platform for retraining and refinement efforts as context-specific data is added or new data becomes available.

Our results demonstrate that FLYNC is not only capable of identifying a significant number of previously unannotated transcripts but also classifies these with high precision and reliability. The application of FLYNC to both bulk and single-cell transcriptomic data illustrates its versatility and effectiveness in discovering novel lncRNAs across different biological contexts and experimental settings. FLYNC’s utility was further evidenced by its application to publicly available transcriptomic datasets, where it successfully identified and classified lncRNAs, providing a foundation for future functional and regulatory studies. Moreover, the integration of SUBCELL with FLYNC extended the pipeline’s applicability to single-cell RNA-seq data, highlighting the potential for cell-type-specific lncRNA discovery.

In summary, FLYNC represents a significant advancement in the study of the non-coding genome of *Drosophila melanogaster*. It is a powerful and adaptable tool that streamlines the discovery of lncRNAs, enabling researchers to uncover the hidden layers of RNA regulation. As such, FLYNC is a valuable addition to the toolkit of geneticists and molecular biologists, promising to enhance our understanding of the complex regulatory networks that underpin biological function and disease.

## DATA AVAILABILITY

FLYNC is an open source software available through an GPL-3.0 license in the GitHub repository (github.com/homemlab/flync).

SUBCELL is an open source software available through an GPL-3.0 license in the GitHub repository (github.com/homemlab/subcell).

## ACKNOWLEDGEMENTS

We thank the cytometry and fly facilities at NOVA Medical School for technical support and CONGENTO: consortium for genetically tractable organisms (LISBOA-01-0145-FEDER-022170), Bloomington *Drosophila* Stock Center (NIH P40OD018537) and Vienna *Drosophila* Resource Center (VDRC) for the stocks used in this study. We thank all members of the C.C.F.H. Lab for helpful discussions. We would also like to thank the National Distributed Computing Infrastructure of Portugal (INCD) for providing the required resources to compute memory-intensive tasks. INCD was funded by Fundação para a Ciência e Tecnologia (FCT) and FEDER under the project 01/SAICT/2016 n° 022153. This project has received funding from the European Research Council (ERC) under the European Union’s Horizon 2020 research and innovation programme (H2020-ERC-2017-STG-GA 759853-StemCellHabitat to C.C.F.H.); by Wellcome Trust and Howard Hughes Medical Institute (HHMI-208581/Z/17/Z-Metabolic Reg SC fate to C.C.F.H.); European Molecular Biology Organization (EMBO) Installation grant (H2020-EMBO-3311/2017/G2017 to C.C.F.H.) and EMBO Long-Term Fellowship (EMBO-ALTF/1208-2018 to T.B.); by FCT (PTDC/BIA-BID/0681/2021 and IF/01265/2014/CP1252/CT0004 to C.C.F.H.; CEECIND/02989/2017 and EXPL/BIA-BID/1394/2021 to T.B.). This work was also supported by iNOVA4Health (UIDB/04462/2020 and UIDP/04462/2020) and LS4FUTURE (LA/P/0087/2020) to C.C.F.H.

